# Novel polymer fixed-target microfluidic platforms with an ultra-thin moisture barrier for serial macromolecular crystallography

**DOI:** 10.1101/2025.07.13.663488

**Authors:** Sankar Raju Narayanasamy, Megan L. Shelby, Chandraki Chatterjee, Jenny Zhou, Samuel Rose, Julian Orlans, Swagatha Ghosh, Anne Marie Cardenas, Sabine Botha, Kevin Gu, Donald Petit, Zhongrui Liu, Francesco Fornasiero, Stella Lisova, Elyse Schriber, Daniel Rosenberg, Thej Tumkur Umanath, Silvia Russi, Brent Segelke, Tonya L. Kuhl, Martin Trebbin, Shibom Basu, Daniele de Sanctis, Matthias Frank

## Abstract

The advent of ultrabright fourth generation X-ray light sources, including X-ray free-electron lasers (XFELs) and diffraction limited synchrotrons, has significantly advanced the field of serial macromolecular protein crystallography (SX). SX experiments demand a continuous supply of fresh microcrystalline sample, ideally while minimizing overall sample consumption. Here, we introduce a novel, robust, and user-friendly polymer film technology that can be assembled in various configurations to encapsulate protein microcrystals and provide sample support for SX. This system provides an efficient hydration barrier over extended durations while maintaining an exceptionally low X-ray background. We have validated this technology by assessing hydration retention under both ambient and ultra-high vacuum conditions, and by evaluating its mechanical stability under XFEL pulses. Furthermore, we have demonstrated the effectiveness of this approach in two room-temperature serial crystallography studies to determine the structure of a 24 kDa Rapid Encystment Phenotype (REP24) protein from *Franciscella tularensis*.

## Introduction

By capturing detailed molecular coordinates, macromolecular crystallography provides deep insights into the structural mechanisms of biological function and disease, information that is vital for advancing drug discovery and the development of targeted therapies. Structure-based drug design is among the many applications of high-resolution structural data of target proteins in complex with bound ligands. These data are essential for rational ligand optimization and the development of drug candidates with improved potency, selectivity, and pharmacokinetic properties. However, the predominant technique for protein structure elucidation, conventional X-ray crystallography, often requires data acquisition under cryogenic conditions to mitigate radiation damage, capturing the protein in a static conformation^1,2^. Serial macromolecular X-ray crystallography (SX) in the form of serial femtosecond X-ray crystallography (SFX) or synchrotron-based serial millisecond X-ray crystallography^3^ (SMX) has emerged as an innovative approach that enables structural analysis at room temperature (RT), thereby preserving functionally relevant conformational states and providing a more physiologically relevant depiction of protein-ligand interactions.^4,5^ More room temperature protein structures have been determined using SX than ever before with the increase in access to ultra-bright X-ray synchrotron light sources^6–11^ and X-ray Free Electron Lasers (XFELs)^12–14^. These 4^th^ generation light sources combined with a serial data acquisition approach allow acquisition of (near) damage free macromolecular structures from proteins in microcrystals and enable time-resolved measurements of structural changes on millisecond timescales and femtosecond timescales at synchrotrons^15–18^ and XFELs^19–22^, respectively.

However, the SX approach requires continuous, rapid replenishment of protein microcrystals^23–25^. Several sample delivery methods including liquid jets^26–31^, viscous jets^32–34^, and fixed targets (FTs)^35–40^ have been developed to accommodate this. Gas-dynamic virtual nozzles (GDVNs) generate liquid jets by injecting a microcrystal slurry suspended in a carrier liquid into the X-ray beam^41^. Liquid jets maintain sample hydration but are prone to several challenges including difficulty in flow rate control^42,43^, clogging^44–46^, X-ray scattering background from carrier fluid^47^, crystal damage during injection, compatibility issues between sample and carrier fluid, and most importantly high sample consumption rates.^48^ Similarly, viscous liquid jets have increased X-ray background from the viscous medium and issues maintaining a constant flow rate^33,34^, making SX a challenge for scarce samples.

In contrast, fixed-target delivery systems support crystal samples with a solid support (e.g., a “chip”) mounted in the X-ray beam path and moved by motors to collect diffraction data from crystals positioned inside or on the FT. Various FT approaches have been developed over the past decade^49–61^, including microgrids, free-standing films, and microfluidic chips. FT methods offer several advantages, including reduced sample consumption, more control over the sample environment, the possibility for *in situ* crystallization, and a high degree of customization for different targets. These advantages eliminate the need for complex systems such as pumps and plumbing^62–64^, which typically increase sample-exchange time. Despite these benefits, FT systems have historically had several limitations^65,66^ such as slow scan speeds and thus a low rate of data acquisition, high background caused by the solid sample support, dimensional constraints, complicated loading procedures, incompatibility with ultra-high vacuum conditions, fragility and susceptibility to beam damage, and imperfect control of sample hydration in dehydrating conditions. A range of FT support materials have been explored, including polymeric materials such as polyimide (Kapton), Mylar, and cyclic olefin copolymer (COC). Carbon-based polymeric materials provide lower X-ray absorption than some micropatterned silicon alternatives and are easier to fabricate, making them more practical for FT use. Microfluidic FTs, made from microfabricated or micropatterned solid matrices, allow for the loading of microliter volumes of crystal slurry or the growth of microcrystals directly within the FT. The integration of microfluidics enables precise control over crystal locations, the ability to create complex *in situ* crystallization conditions, adaptation for membrane protein crystallization via Lipidic Cubic Phase (LCP) matrices, and high-precision flow delivery. Despite these advances, combining low-background polymeric films^38,49,56,59,67,68^, simple fabrication, and easy sample loading have remained challenges. These factors have limited the widespread adoption of FT systems as the workhorse sample delivery method for serial crystallography.

To overcome these obstacles, all-polymer fixed-target platform development has become an increased focus of the authors and others^38,39,56,59,65,69^. Here we present a new approach in FT sample delivery utilizing parylene-N (PaN) polymer films that have been engineered to be ultra-thin and highly durable. PaN films demonstrate several favorable properties^70–72^ such as biocompatibility, water impermeability, chemical inertness, a high melting point, structural integrity through higher elongation at break than other X-ray compatible polymers such as Mylar, cyclic olefin copolymer, and Kapton. PaN polymer films are crystalline^73^ in nature, similar to other commonly used X-ray windows such as Kapton. These films are composed of light elements and thus interact minimally with X-rays. Similar to thin films like Mylar, PaN films offer high adaptability for various experimental conditions, and are easy to fabricate, cost-effective, and user-friendly. The PaN FT platform described is robust, well-suited for SX structure determination, and ideal for screening crystals under diverse conditions while minimizing sample usage.

## Results and Discussion

### PaN film characterization and incorporation into microfluidic FT devices

A critical criterion for a FT sample support is a minimal contribution to background during protein microcrystal diffraction data collection, which is essential for achieving high-quality data at atomic resolution. To meet this requirement, 6-inch diameter free-standing, hydrocarbon-based PaN films were fabricated and subsequently cut to size based on beamtime needs (see Methods and Supplementary Material). The fabrication process has been optimized to result in less than +/- 5% overall thickness variation across the 6-inch diameter wafer, corresponding to a maximum thickness variation of less than 200 nm, as shown by the measurement of film thickness using thin-film spot mapping system (Fig. 1a). Given that up to 28 one-inch squares can be cut from a 6-inch diameter film, the thickness variation per individual square is limited to approximately 20–30 nm, ensuring high uniformity and consistency across individual samples. To demonstrate the robustness of such PaN films, a 1.2 µm thick PaN film sample was clamped between two copper gasket O-rings using paper clips, creating an open, free-standing film area inside the rings with a diameter of 37 mm. A weight of 400 grams was placed on top and the film remained intact, showing no signs of mechanical damage (Fig. 1b). Unlike Mylar films of similar thickness, which are susceptible to breakage and therefore difficult to handle, the PaN films exhibit excellent mechanical resilience and ease of handling.

**Figure 1:**
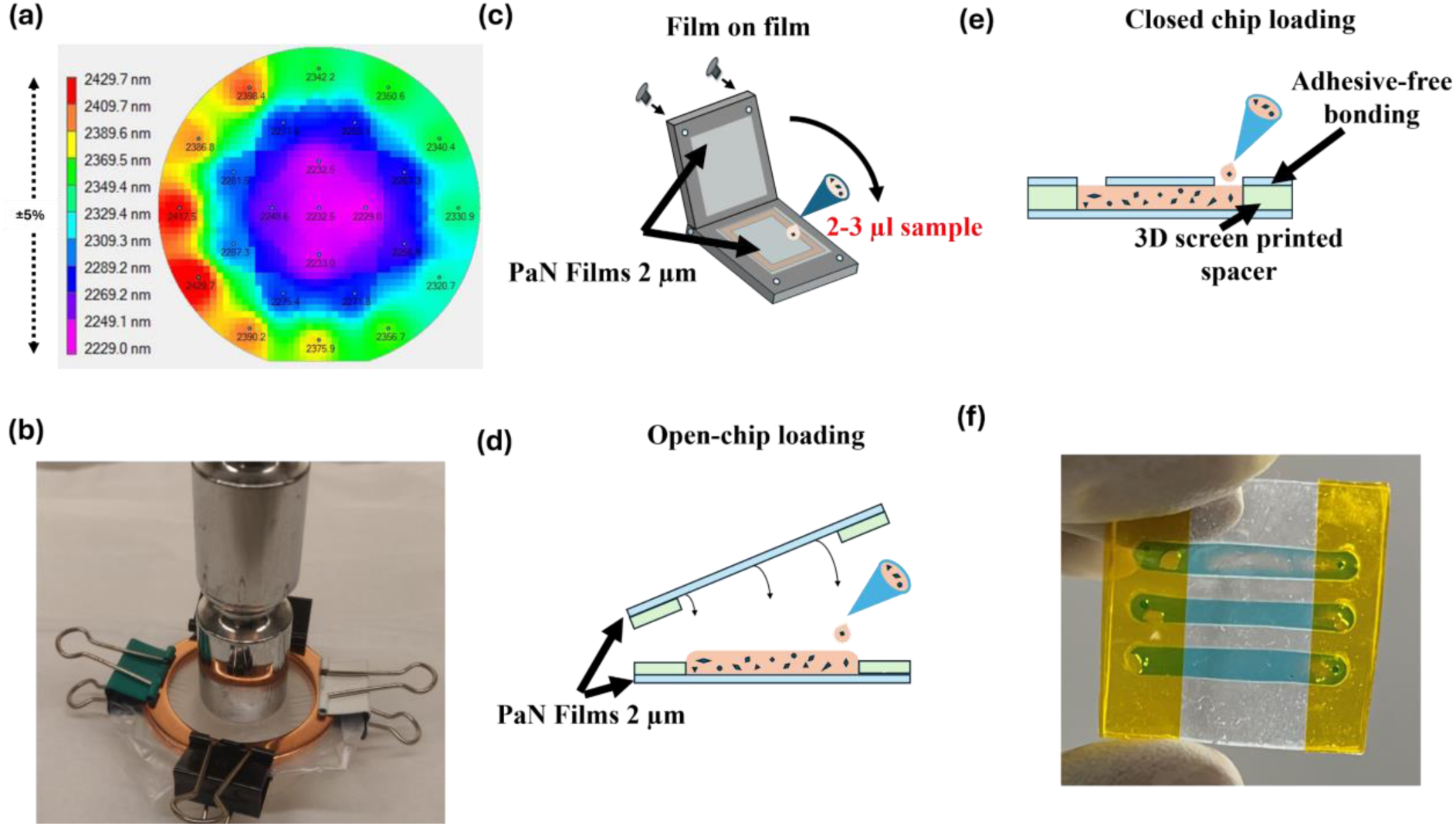
Characterization of PaN films and incorporation into microfluidic devices and chips: a) Thickness variation map of a ∼2.3 μm thick free-standing Parylene-N (PaN) film measured using Filmetrics-Thin-Film Small-spot Mapping system depicting a thickness variation across the 6” wafer of only +/- 5%.; b) Structural Integrity of a 1.2 µm PaN film with 37 mm diameter demonstrated by placing a 400 g weight on it; c) Schematic of the film-on-film sample loading-PaN fixed-target platform. Sample is placed on the bottom PaN film and then sandwiched with the top PaN film and held in place mechanically in between O-rings and a metal holder. This FT is ideal for ambient condition experiments with sensitive crystal samples that are fragile, and for viscous sample; d) Schematic of the open-chip loading fixed-target platform. e) Schematic of closed-chip fixed-target platform with separate, independent windows. Here the sample is loaded by pipetting through loading channels in the top layer of the PaN FT and is drawn into the microfluidic channel using capillary force; f) A fully assembled closed-chip loading fixed-target platform loaded with blue color dye solution.

**Figure 2:**
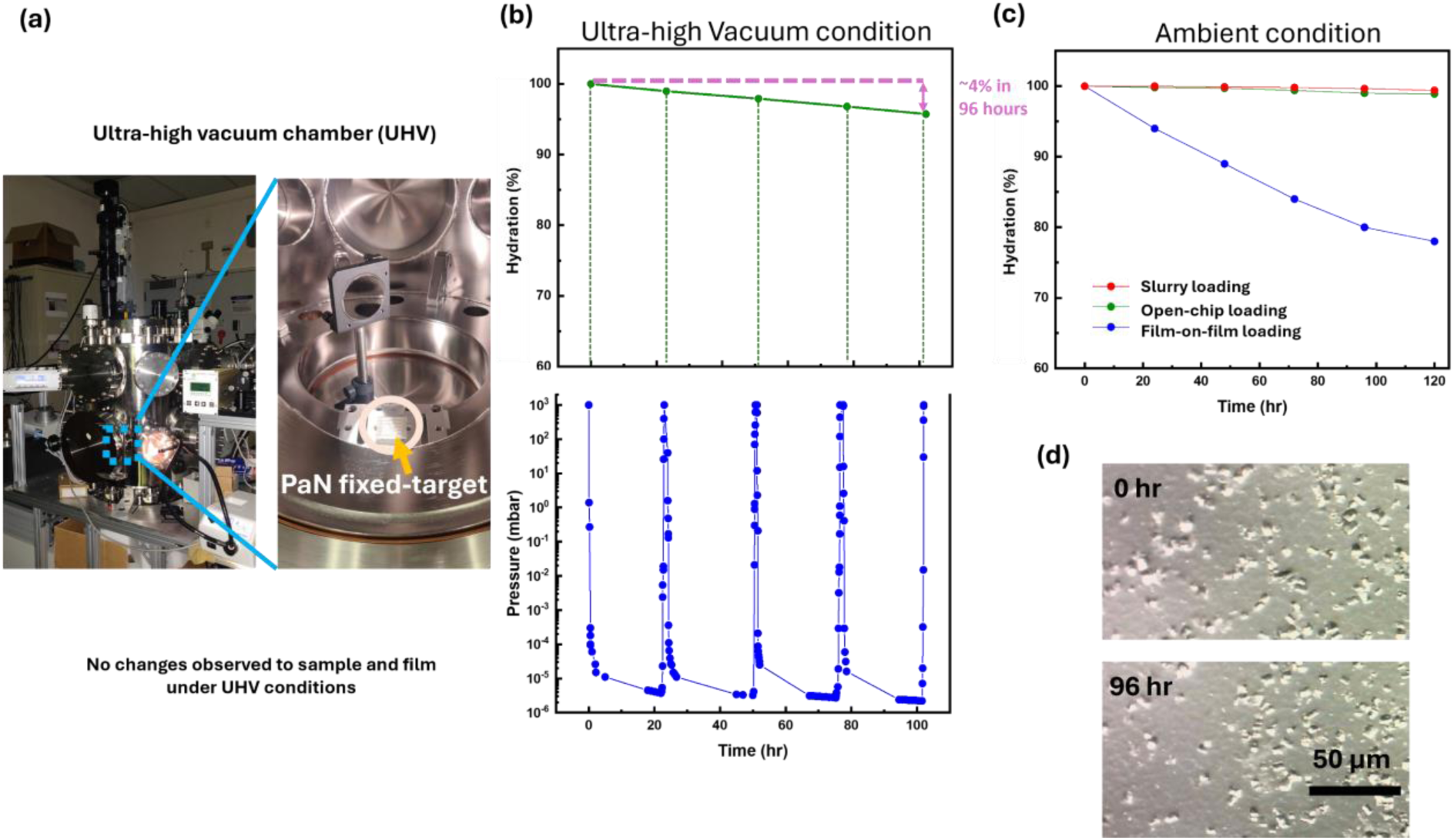
Hydration retention study in PaN fixed-target platforms: a) Ultra-high vacuum UHV) chamber. An inline camera was placed inside the chamber to visualize any damage to the ultra-thin PaN films during any of the pump-down and vent cycle. No damage to the PaN films was observed despite multiple UHV cycles.; b) Hydration retention study of PaN fixed-target platform loaded with a 1:1 ratio of water and REP24 crystallization buffer (final concentrations 27%(v/v) PEG-MME 750, 50 mM Na-acetate pH 4.5 in 25 mM NaCl, and 5 mM HEPES pH 7.5) in UHV chamber. The mass of the PaN fixed-target platform was measured after each 24 hour cycle. The results show that the hydration loss is less than 4% over 108 hours under ultra-high vacuum conditions; c) Hydration retention study of REP24 protein crystals in ambient conditions compared under loading various PaN FT platform types, such as film-on-film loading, open-chip loading, and closed-chip loading. The results show that for a mechanically clamped setup in film-on-film loading, there is a loss of ∼20% over a period of 120 hours, and we hypothesize that this loss is through the sides held by a rubber O-ring in the SOS chip holder^49^ and not loss through the PaN films. And for closed-chip PaN FT crystal slurry loading and open-chip loading PaN FT there is near to no significant observable loss of moisture since they are completely sealed on the sides by the PDMS screen printed structure, and the PaN film eliminated moisture loss in the window region; d) The crystals undergo no apparent damage after 96 hours under UHV conditions..

**Figure 3:**
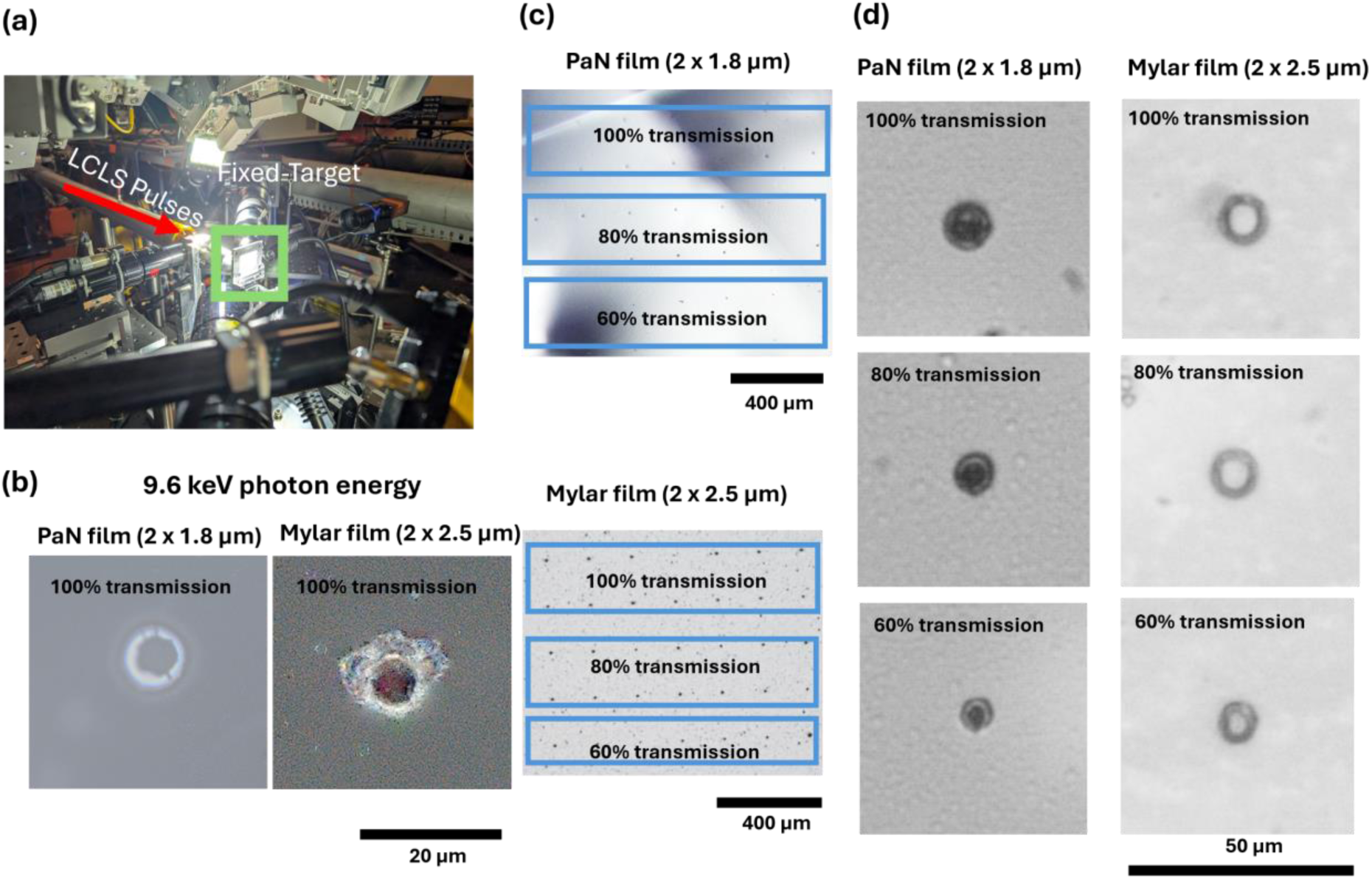
XFEL damage study on thin films used for fixed-target platforms: a) Experimental setup with PaN fixed-target platform at the MFX beamline at the LCLS; b) Comparison of film damage caused by single XFEL pulses observed by z-stack extended focus image of PaN film (2×1.8 µm thickness) and Mylar film (2×2.5 µm thickness). It is observed that at 100% X-ray transmission the PaN film undergo indentation or dent; c) PaN and Mylar® films investigated at x-ray transmissions from 60 % to 100 % with a shot spacing of 200 µm; d) Comparison of film damages between PaN and Mylar films at X-ray transmissions from 60 % to 100 %. Comparing the damage to PaN film and Mylar film at the same transmissions it is clearly observed that PaN films show minimal visible damage in form of dents, thus making it an ideal polymer film for making fixed-targets.

PaN films were used to construct three distinct types of FT sample support platforms, tailored to different crystal sizes and beamline conditions such as ambient pressure or ultra-high vacuum environments. The first type, a film-on-film FT platform (Fig. 1c), consists of two PaN films sandwiched together with microcrystalline sample suspended in between and mechanically secured with metal clamps containing O-rings. This type of platform was previously demonstrated using thicker Mylar films and was termed as “sheet on sheet” or “SOS”^50^. This configuration is suited for ambient-condition SX experiments and accommodates both aqueous and viscous samples. The second type, an open-chip loading platform, includes a thin polymer spacer between the PaN films (Fig. 1d) that defines the thickness of the sample compartment(s), see Supplementary Material for spacer thickness options. After sample loading onto the open chip (consisting of the bottom PaN layer and the spacer) the top PaN layer is placed on top, and the sides of the chip are sealed by the strong, covalent siloxane bond formed on contact by the Polydimethylsiloxane (PDMS) spacers (see Methods). Such an open-chip platform can be used for viscous or sensitive samples and was employed in this work to support larger crystals in specific chip locations for rotation-series-based serial crystallography^38^. The third loading type is on a closed-chip PaN FT, which is a pre-sealed chip with a thin polymer spacer between the PaN films that includes individual, independent sample compartments with sample loading channels (Fig. 1e, f) and can hold multiple samples at the same time. The closed-chip FT is fully assembled and sealed around the edges before the sample loading. Sample is pipetted into the designated loading ports that are also sealed after the sample loading. The closed chip FT is compatible with SMX experiments that use relatively low viscosity samples. Both open-chip and closed-chip formats are demonstrated here for experiments conducted in ambient conditions, and measurements of their water retention in vacuum suggest compatibility with crystallography in ultra-high vacuum conditions.

### Hydration retention study with PaN fixed-target platforms

Hydration preservation is key for any measurement of protein samples at room temperature, be it in vacuum, or air or helium environment, because loss of hydration can lead to rapid sample degradation.

FT sample supports therefore are required to be robust and maintain structural integrity under X-ray irradiation. Of the three environments, a vacuum environment contributes the least to X-ray scatter background, making it ideal for serial crystallography studies, in particular, for weakly scattering nano-or microcrystals but poses challenges for sealing the sample (compared to, e.g., a humidified helium environment).

Another key development is the compatibility of open-chip loading PaN FT, and closed-chip PaN FT with ultra-high vacuum (UHV) environments for extended periods of time. Synchrotron and XFEL experiments often require ultra-high vacuum conditions^74^ to prevent air scattering of the X-ray beam. Our polymer film-based platform is specifically tailored to be ultra-high vacuum compatible, ensuring that the samples remain stable and intact throughout the experiment. In addition to its vacuum compatibility, the polymer film also plays a crucial role in preventing hydration loss from the protein crystals during data collection, and allows for storage of samples for long durations.

Our hydration retention studies conducted on PaN FT platforms aimed to evaluate the moisture loss and overall stability of the platform in UHV and ambient conditions. In the UHV study, a closed-chip PaN FT, with a film thickness of 2.4 µm was loaded with 4 µL of buffer (each chip had 2 films, i.e., 2 × 2.4 µm), its loading ports sealed, weighed and then placed in an UHV chamber that was pumped down to <10^-5^ mbar. The PaN FT platform was held in vacuum for a total of ∼4 days. During that time the chamber was briefly vented each day to measure the loss of moisture through the windows by weighing the fixed-target platform (with the reduction in weight being attributed to water loss), and then introducing it again into the UHV chamber. To observe any potential damage of ultra-thin PaN films during pump-down and vent cycles, an inline camera was placed. Despite multiple UHV cycles, the PaN films remained visibly intact and free from any damage, demonstrating their robustness and reliability under vacuum conditions. The results indicated that the moisture loss during the 108-hour study was less than 4% from overall volume of sample loaded, suggesting that the PaN platforms are highly effective at maintaining hydration even under the extreme conditions of ultra-high vacuum. We repeated the same with REP24 protein crystal slurry loaded sample in closed-chip PaN FT, and with open-chip loading PaN FT and the results were the same.

We performed another hydration retention study in ambient conditions. In this study, we investigated the loss of hydration of REP24 protein crystal samples, comparing different types of PaN fixed-target platforms, including film-on-film loading, open-chip loading, and slurry loading. The results showed that with the mechanically clamped setup in film-on-film loading, there was a significant water loss of about 20% over 120 hours. This loss was hypothesized to occur through the sides held by a rubber O-ring in the SOS chip holder^50^ rather than through the PaN films. In contrast, for slurry loading and open-chip loading platforms, which were completely sealed on the sides, there was virtually no water loss (<1%), indicating that the PaN film effectively prevented water loss in the window region, and the protein crystals experience no damage, confirming that the PaN platform not only preserved hydration but also maintained the integrity of the crystals over extended periods. These hydration retention studies demonstrate that PaN FT platforms are highly effective in minimizing water loss, particularly in ultra-high vacuum conditions, while also maintaining the structural integrity of both the platform and the protein crystals.

### XFEL Damage Study on a PaN FT platforms

The study was conducted at the Macromolecular Femtosecond Crystallography (MFX) beamline at the Linac Coherent Light Source (LCLS) operating at a photon energy of 9.6 keV, X-ray pulse energy of ∼1.6 mJ (at 100 % transmission), X-ray spot size estimated to be ∼3 µm, and a repetition rate of 120 Hz. Both PaN films in a PaN fixed-target platform and and Mylar films in a film-on-film FT platform were exposed to X-ray pulses at different X-ray transmission levels. Multiple shots were taken on the same films, spaced 200 µm apart, to assess the cumulative effect of XFEL pulses on the films’ structural integrity.

These images show that, even at 100 % transmission, the PaN films exhibited only localized indentation or denting, while the Mylar films sustained more severe damage, including hole formation. In these experiments, damage to the films by X-ray pulses is likely caused by X-ray absorption and the resulting heating and thermal shock of the films and is expected to depend also on the film’s melting points and elongation at break. At 9.6 keV, PaN has about a factor 2 lower X-ray absorption (∼3750 µm attenuation length) than Mylar (∼1860 µm attenuation length), and thus for the same film thicknesses about 2x more energy is expected to be deposited in Mylar. (Note that the Mylar films used in these experiments were slightly thicker than the PaN films.) In addition, PaN has higher elongation at break than Mylar (200% for PaN, and ∼70% for Mylar) and a higher melting point than Mylar (420^0^C for PaN and 254^0^C for Mylar). This damage study highlights the superior durability of the PaN film under high-fluence conditions compared to Mylar.

### Room temperature structure determination of REP24 using serial crystallography

These PaN-based FT platforms have also been successfully deployed for structural studies at several synchrotron and XFEL facilities (Fig. 4). At ESRF and SSRL, they were used to determine the structure of REP24, a putative virulence factor from *Franciscella tularensis*.^75^ The PaN fixed-target platforms were loaded with REP24 protein crystals (Fig. 4a,b) and were studied at room temperature at Beamline 12-1 in the Stanford Synchrotron Radiation Light source (SSRL), and at Beamline ID-19 in the European Synchrotron Radiation Facility (ESRF).

**Figure 4:**
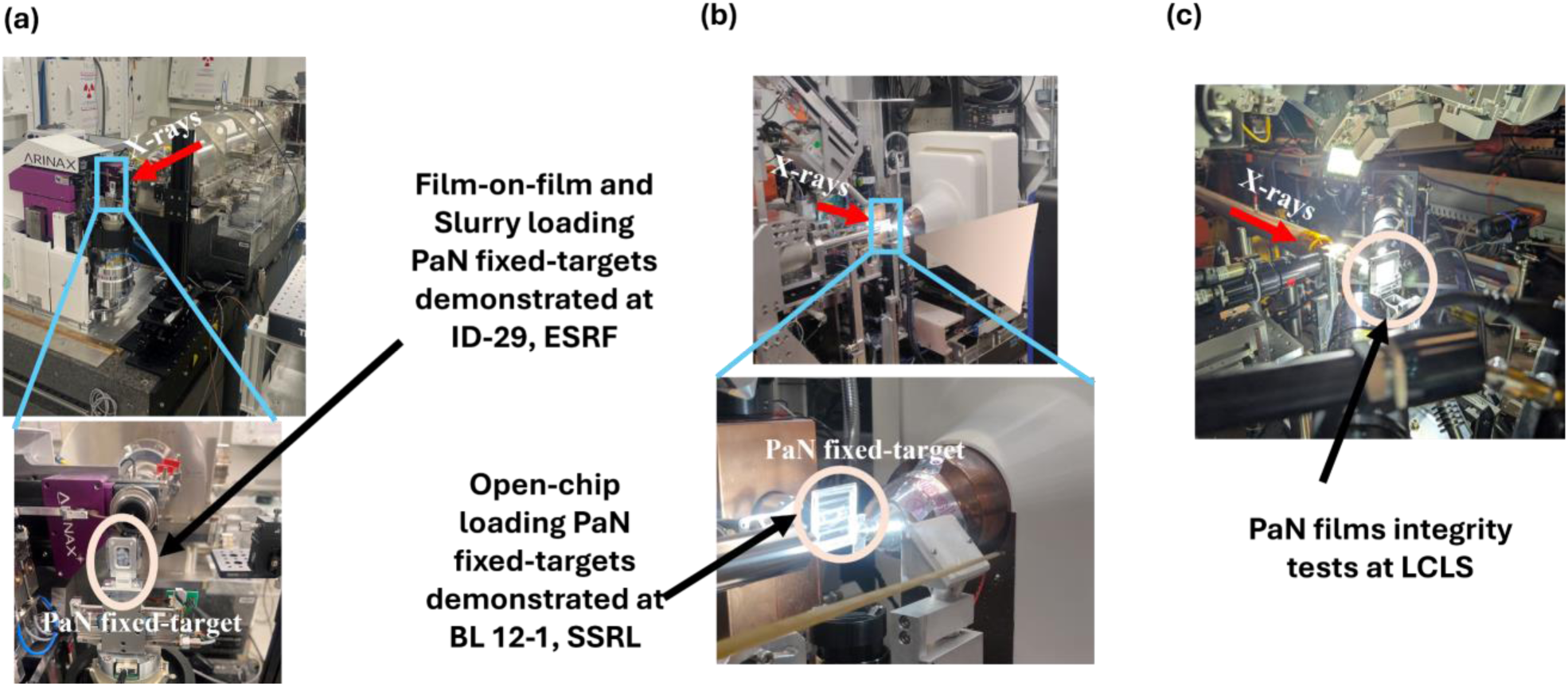
PaN Fixed-target platforms implementation in beamlines. a) Demonstration of PaN fixed-target platform using both film-on-film loading PaN FT and closed chip PaN FT at ID-29 Serial Macromolecular Crystallography beamline at ESRF. Both FTs were held in place to the actuators using SOS metal holders^49^; b) Demonstration of open-chip loading PaN FT with large REP24 crystals (∼100 µm) at SSRL, SLAC; c) Film on film PaN FT were introduced to the X-ray Free-Electron Laser (XFEL) beam at LCLS, SLAC.

The PaN films exhibited excellent x-ray transparency (Fig.5) without compromising mechanical stability and hydration retention during the duration of the measurement. At SSRL, short rotation series at 12.4 keV were collected for several individual crystals and merged to form a complete dataset. This strategy utilized a crystallization condition producing large, approximately 100 um crystals of REP24 applied using the open-chip PaN FT. At ESRF, data was collected at 11.6 keV using a raster scanning approach utilizing a high crystal density. Crystals were loaded using the film-on-film PaN FTs. The crystal sizes were in the range of 15-20 µm. In both cases, no evidence of crystal degradation or dehydration during the duration of measurement was noted.

**Figure 5:**
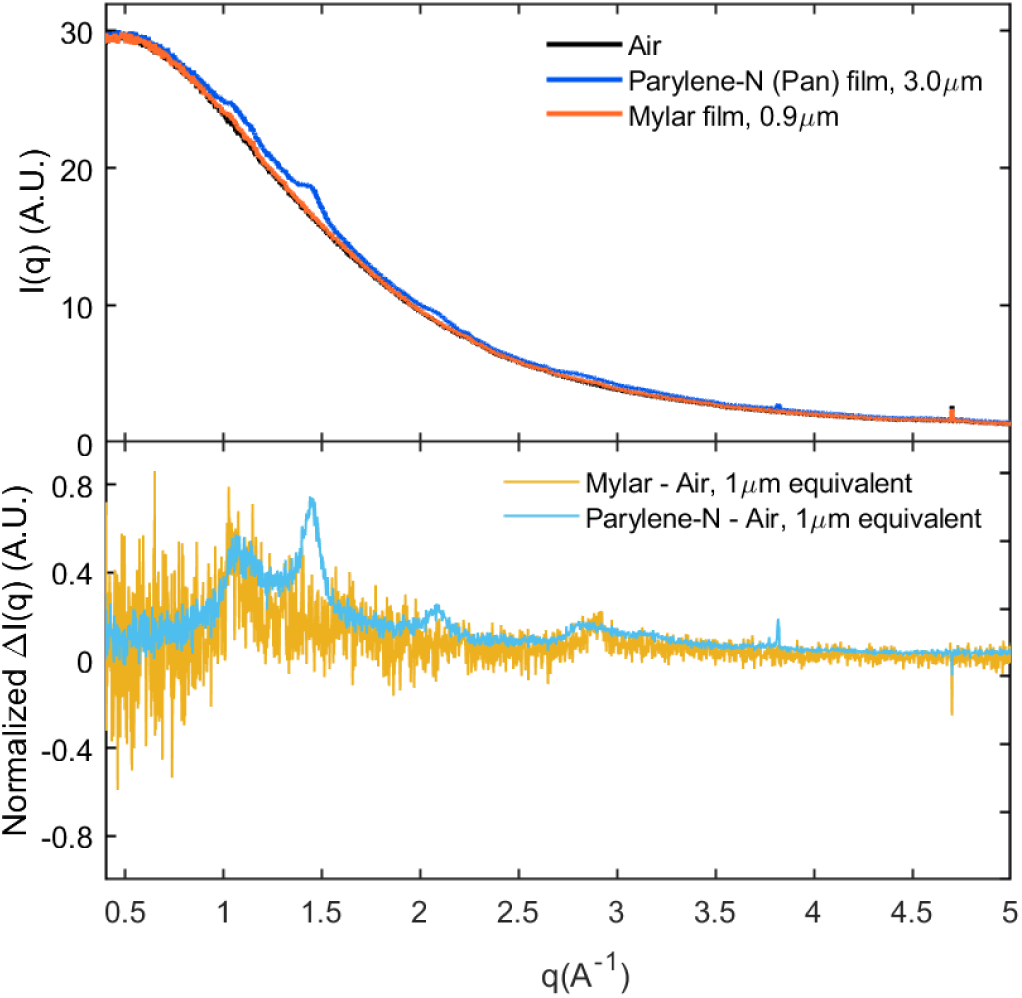
X-ray transparency of PaN films: a) A comparison of radial averages of film and air scattering at Beamline BL 12-1, SSRL demonstrate that the background contributed by non-crystal and non-buffer experimental components is dominated by air scattering at 12.4 keV. Scattering observed from the thinnest available Mylar, and PAN films are comparable to air alone. b) The air-subtracted film scattering, normalized by film thickness, is similar in magnitude at q=1 and 3, though PAN displays several more well-defined peaks at q=1.5,2.1.

**Figure 6:**
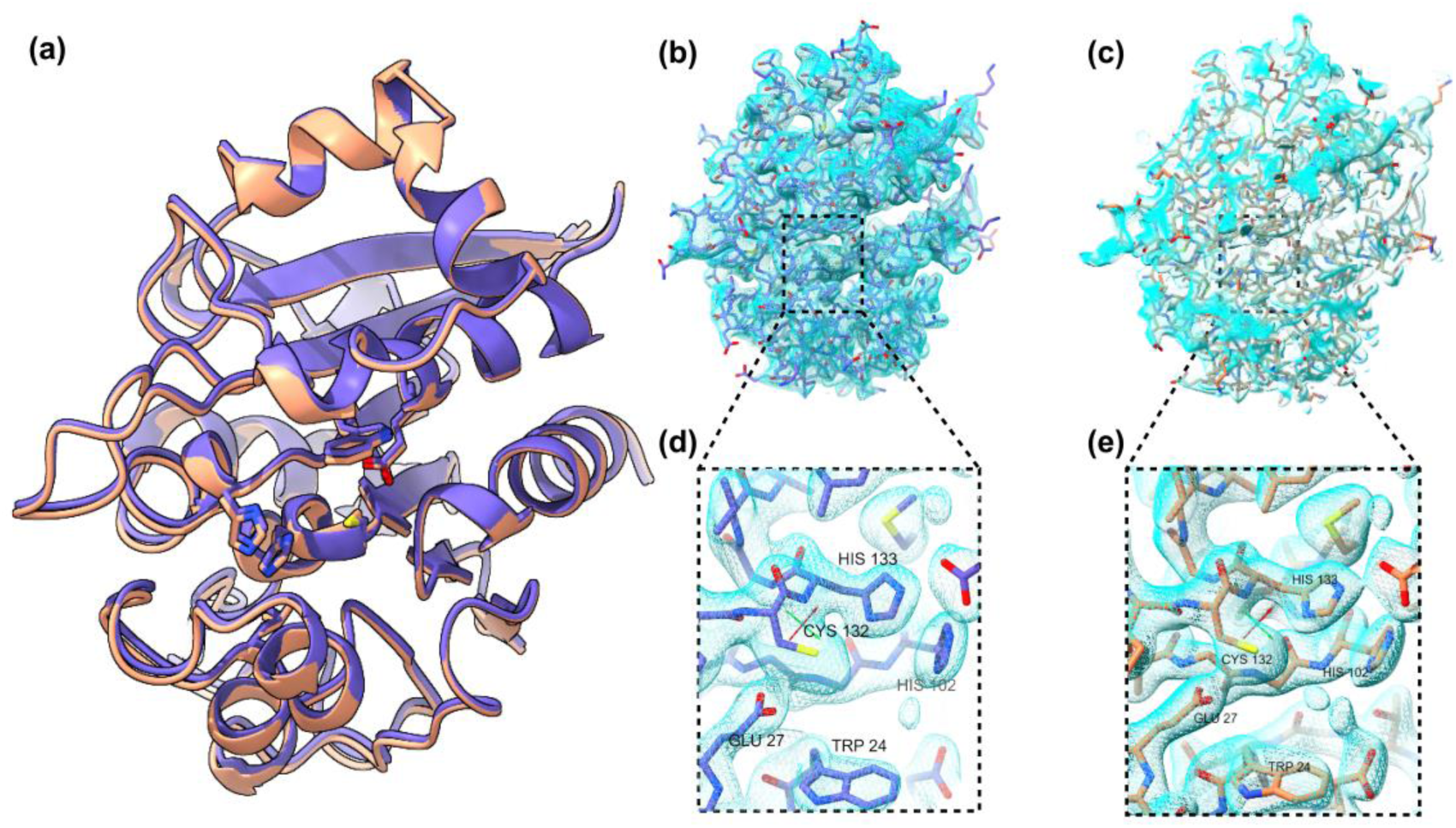
a) Overlaid structures of Rapid Encystment Phenotype Protein 24 kDa (REP24) obtained using merged rotational datasets from multiple crystals at room temperature at the Stanford Synchrotron Radiation Source (SSRL) beamline 12-1 (purple) and synchrotron serial crystallography at the European Synchrotron Radiation Facility (ESRF) (peach). In both cases REP24 was enclosed in PAN thin films during data collection. Sidechains of putative active site residues W24, E27, H102, C132, and H133 are shown. Electron density maps were reconstructed to 1.6 Å and 1.8 Å at b,d) SSRL and c,e) ESRF respectively. Models were refined from an existing structure solved using cryo single-crystal crystallography (4P5P)

Electron density maps reconstructed from both datasets are similar in quality, though slightly higher resolution data was obtained from SSRL serial data collection. Models were refined using the deposited single crystal structure of REP24 measured in cryogenic conditions (4P5P) as an initial model. Sidechains of putative active site residues W24, E27, H102, C132, and H133 are well resolved in both structures.

## Conclusions

Serial crystallography (SX) at room temperature enables high-resolution structural studies of proteins in their biologically relevant states. To address challenges in conventional delivery methods, such as high sample consumption, background scattering, and limited vacuum compatibility, we developed fixed-target (FT) platforms using ultra-thin parylene-N (PaN) films. PaN combines low X-ray absorption, exceptional mechanical resilience (elongation at break ∼200 %), high thermal stability (melting point ∼420 °C), and effective moisture retention. These films are easy to fabricate, cost-effective, and user-friendly, offering rapid assembly and flexible sample loading without complex plumbing or pumps. PaN FTs preserve hydration in both ambient and ultra-high vacuum (∼10⁻⁵ mbar) environments, with less than 4% water loss over 108 hours shown through an example. Under XFEL exposure (9.5 keV, ∼2.26 × 10⁴ J/cm² fluence), PaN films only exhibited denting, while Mylar films were perforated, highlighting PaN’s superior durability. PaN films are highly resilient to XFEL-induced damage and exhibit excellent moisture retention, making them an ideal choice for fixed-target platforms where both durability and hydration control are critical. This dual functionality not only ensures greater stability for sensitive samples but also highlights the potential of PaN FT platforms for advancing serial crystallography studies at XFELs. Furthermore, wafer-scale deposition of PaN films, and roll-to-roll fabrication of PaN FTs enables us to produce sufficient quantity of fixed-target platforms at ease and on-demand based on beamtime requirement. PaN FTs were successfully used for room-temperature structure determination of REP24 at SSRL and ESRF, confirming their utility as robust, low-background, and vacuum-compatible platforms for high-throughput serial crystallography, opening new avenues for structural biology research for a broader range of proteins, particularly at synchrotron and XFEL facilities.

## Methods and Development

### Fabrication of free-standing PaN films

The free-standing PaN films are prepared by Parylene-N dimer, a powdery raw material (Specialty Coating Systems, Inc.). The Parylene-N is poly(para-xylylene), a completely linear, crystalline material with high dielectric strength, biocompatible, excellent moisture and chemical barrier properties, UV stable, and highly conformal to the required shape. The initiated chemical vapor deposition (iCVD) was utilized to deposit the Parylene dimer on the Silicon wafers. The wafers were lined up in the chamber and the pressure and temperature were set to the chamber size and appropriate quantity of dimer was introduced to form a thin layer on the silicon wafers. To reduce thickness variation below 20%, we have introduced custom stage at middle of the chamber. Using this customized chemical vapor deposition, we can obtain thin free-standing PaN films, as thin as 27 nm. For ease of film removal from the silicon wafers, we tested several lift-off methods. We used sacrificial layers for lift-off, some prominent ones are polyvinyl alcohol (PVA), glycerine, micro-90 soap solutions, and a composite method of combining with ultra-thin PET film (900 nm). For our SX studies that require batch fabrication of these large ultra-thin PaN films of constant size, we utilized diluted micro-90 soap solution as sacrificial layer. After the lift-off, each film is rinsed by placing it in a sonicator filled with iso-propyl alcohol for 20 minutes to remove any remaining soap solution or other dust. Then these films are carefully transferred to cleanroom wipes and stored for future use. Unlike thin COC-films that degrade over time, these PaN films have shown no signs of degradation even after a period of 24 months. Our methodology of placing the wafers precisely at the center of the deposition chamber resulted in extremely uniform PaN films, with variation of less than 200 nm (+/- 5%) over the entire 6-inch wafer. Through our batch fabrication we made 2 µm thick PaN films for our beamtime experiments.

### Assembly of Film-on-film PaN fixed-target platform and loading sample

The large free-standing PaN films are simply cut to size using a razor blade, and two such films are used to prepare a film-on-film PaN fixed-target platform. The first film is placed on one side of the SOS holder^50^, then ∼3-5 µl of crystal slurry is placed as small droplets on the film and the second film is placed on top of it. Then, the other side of the metal holder is placed upon the two films, and mechanically held in placed with screws. This methodology is suitable for all ambient condition SX studies.

### Assembly of Open-chip PaN fixed-target platform and loading sample

Open-channel chips are desired for large microcrystals that need gentle handling but can be utilized for others as well. For assembling the open-chip loading version (Fig. 1d), we plasma treat two halves of the fixed-target platform, then we place the bottom half on a flat surface, load the crystals gently using large pipette tip, and then place the other half on the top, the top half usually is supported by a thin layer of PMMA or Aluminum frame (500 µm) for easier handling. After sealing the fixed-target platform, we apply a thin layer of clear nail polish (Ted Pella Inc.) on the sides to increase the sturdiness of the chip. The nail polish step can be skipped if utilized in room temperature conditions but highly suggested for UHV chamber experiments. Crystals grown in wells by batch crystallization were collected and placed on the PaN sheet (lower bottom) and was sandwiched with another half of PaN sheet (top), both sheets were held by thin PMMA frames (500 µm) for ease of handling.

### Assembly of Slurry loading PaN fixed-target platform and loading sample

When chips are desired for ultra-high vacuum conditions or potential automation of FTs, the closed-chip PaN FT would be preferred. To fabricate this platform, two 2×3 cm pieces of ultra-thin PaN films are prepared. Each piece is screen printed with a layer of PDMS and baked at 100°C. The PDMS-coated surfaces are then treated with plasma, brought into contact, and bonded under mechanical pressure, resulting in robust, adhesive free bonding. The merged PDMS serves as the spacer layer. The spacer thickness can be varied between 10-80 µm, on demand per the crystal size range. The top film is perforated with ∼1 mm holes using a femtosecond laser or mechanical punch. The films can be cut to desired shapes and sizes using femtosecond laser^76^. During the beamtime, based on the sample holders desired at the beamlines, the films can be kept as it is in the SOS metal holders, or plastic holders or can be attached to thin PMMA frames, thus making it a versatile version. The sample is then slurry loaded by pipetting through the hole in the top film, with the liquid filling the internal cavity through capillary action.

### Hydration retention study methodology for PaN fixed-target platforms

The Hydration retention studies of PaN fixed-target platforms were performed in both ultra-high vacuum conditions and ambient conditions over a period of 108 hours and 120 hours, respectively. A UHV chamber that pumps down to 10^-6^ mbar was utilized, and the mass of the fixed-target platform was measured using a weighing scale with a sensitivity of 1 mg. For UHV based dehydration study, the chamber pumped down and vented every 24 hours, and mass of the PaN fixed-target platform was measured during each cycle. For ambient condition study, the chip was left as it is in the ambient condition and the mass was measured every few hours. The percentage of weight loss in the fixed-target platform was calculated as the change in weight loss (attributed entirely to loss of water) over the weight of the liquid sample loaded at the start.

### Expression, purification, and crystallization of REP24

REP24 [24 kDa; PDB code 4p5p^77^], a putative virulence factor protein from the intracellular pathogen Francisella tularensis, was expressed recombinantly and purified as described previously.^35,78^ Crystals for SSRL experiments were grown in vapor diffusion conditions using an 8 µl drop and 800 µl reservoir volume. The drop consisted of a 1:1 mixture of 14.4 mg ml^−1^ REP24 sample (in 50 mM NaCl and 10 mM HEPES pH 7.5) and a precipitation solution (also used in the reservoir) containing 27%(v/v) PEG-MME 750 and 100 mM Na-acetate pH 4.8.

Batch crystallization conditions were utilized for the production of the REP24 crystals in bulk for ESRF experiments. REP24 crystals were grown by mixing a 14.4 mg ml^−1^ REP24 sample (in 50 mM NaCl and 10 mM HEPES pH 7.5) with a precipitation solution containing 25%(v/v) PEG-MME 750 and 100 mM Na-acetate pH 5.5 in a 1:1 ratio. The crystals of REP24 used in the experiment were between 15 and 20 µm in length and had the appearance of two square-based pyramids connected at the peak, with a maximum thickness of 10 µm. Crystal concentrations were estimated at 2.2 × 10^6^ crystals ml^−1^ based on counts done via optical microscopy of a fixed volume.

### XFEL damage study methodology

To understand the damage on PaN films by XFEL beam, we performed the beam damage study at Macromolecular Femtosecond Crystallography (MFX) beamline in Linac Coherent Light Source (LCLS), at SLAC National Accelerator Laboratory. We placed two 1.8 µm PaN films on SOS metal holder^50^ and introduced to X-rays of 9.6 keV photon energy (1.6 mJ pulse energy at 100 % transmission), with X-ray spot size estimated to be ∼3 µm, and a repetition rate of 120 Hz. At 100 % X-ray transmission the X-ray fluence was calculated to be 2.26 × 10⁴ J/cm² (equivalent to 1.47 × 10¹⁹ photons/cm²; see calculations in the supplementary materials), and were tested for 60%, 80%, and 100% X-ray transmission. Multiple shots were taken on the same films, spaced 200 µm apart, to assess the cumulative effect of XFEL pulses on the films’ structural integrity. The films were then visualized with Keyence VHX-7000 microscope with maximum intensity projection technique to study the damage caused by XFEL beam on the ultrathin PaN films. And a comparison was done with Mylar films (2x 2.5 µm thickness).

### Serial macromolecular crystallography data collection methodology by rotation series at SSRL

A rotation series serial crystallography method was followed for data collection at SSRL. Considering the beam size of 5 × 40 µm at the 12-1 beamline in SSRL, larger crystals (∼100 µm) were utilized, and the samples were housed in a open-chip loading PaN-fixed target platform. The assembled chip with REP24 crystals was clipped to a standard magnetic mount, which was in turn secured onto a slotted holder with a magnetic base using a screw, and this assembly was subsequently mounted onto the goniometer at the beamline. Diffraction data were collected at a wavelength of 0.9799 Å, using an Eiger X 16M detector (Dectris AG) at a detector distance of 0.2 m, at 11.56 keV photon energy. The beamline’s sample holder translational motors were used to align and center individual crystals in the beam path, using the inline high-resolution camera to identify each crystal. Datasets were collected from these centered single crystals in 30° rotations. Diffraction data nine sweeps were merged to complete datasets for REP24 with xia2^79–82^, and the structure were refined with *Phenix 1.19*^83^.

### Serial macromolecular crystallography Data collection methodology by raster scanning at ESRF

The raster scanning method of serial crystallography was performed in ambient conditions at the serial macromolecular crystallography beamline ID-29, ESRF-The European Synchrotron. REP24 crystals were concentrated by settling and were housed in both film-on-film loaded PaN fixed-target platform, and in closed-chip loaded PaN FT platform. In both cases, the SOS metal frame was used to hold the PaN FTs, to make ease for sample handling through the beamtime, a 15×13 mm window was fabricated for closed-chip PaN FT. A total of 3-5 µl of crystal slurry was used for each fixed-target platform. From each run we obtained more than 81800 images, and in about five to six runs on the same chip we collected over 561022 images. Each run on the same chip was performed by changing the X-ray shot spacing, thus hitting a different part of the chip each time. 4 × 2 µm (H × V) beam of 11.56 keV photon energy of X-rays, with a 90 µs pulse and 231.25 Hz repetition rate was utilized. Shot spacing of 15-25 µm was chosen for each run. A JUNGFRAU 4M detector with sample-to-detector distance of 103 mm (1.6 Å in the corner) was used to collect diffraction patterns. This technique allowed us to obtain a complete structure of the sample with only one chip, i.e., with only ∼5 µl of crystal slurry.

**Table 1:**
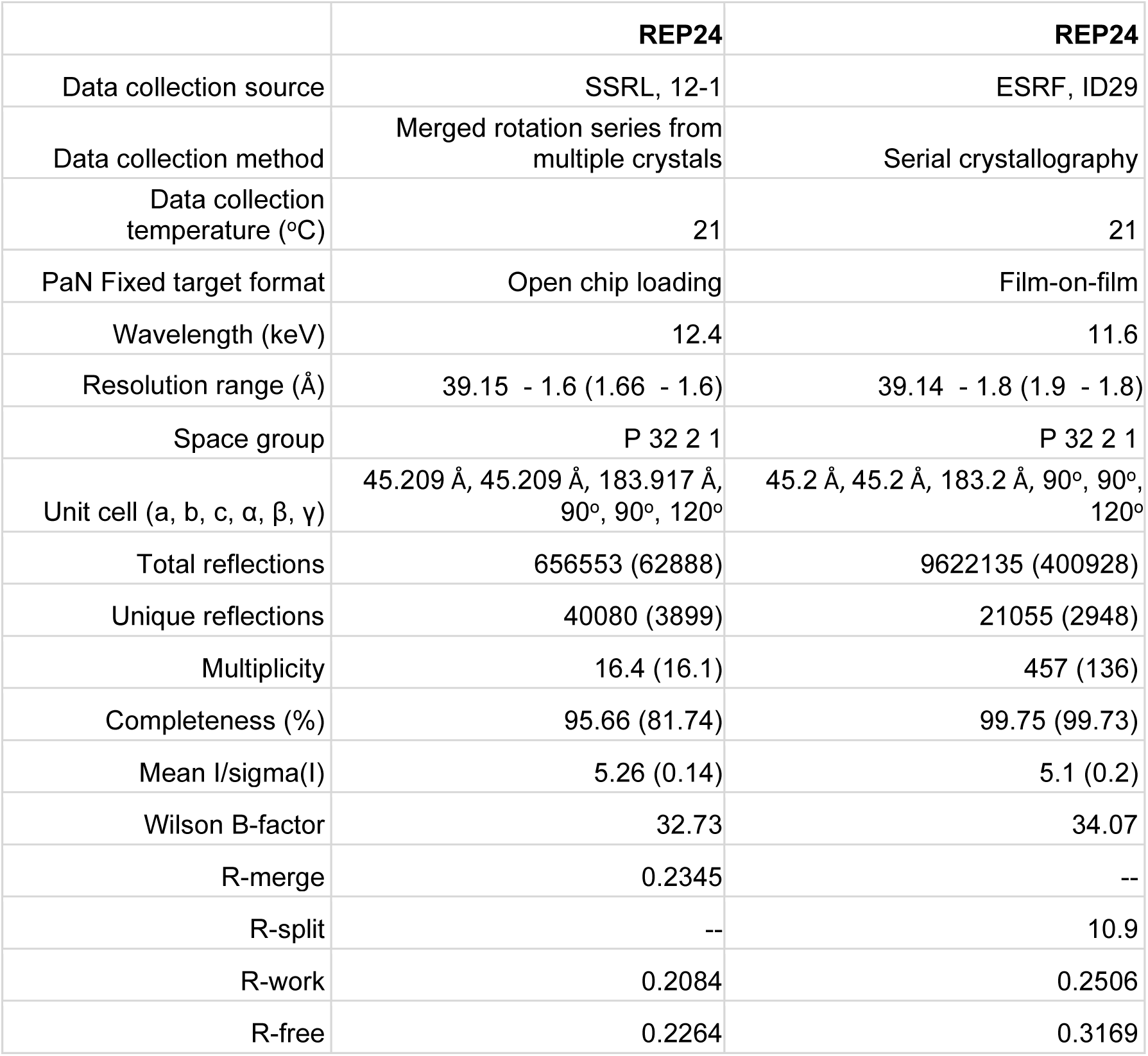
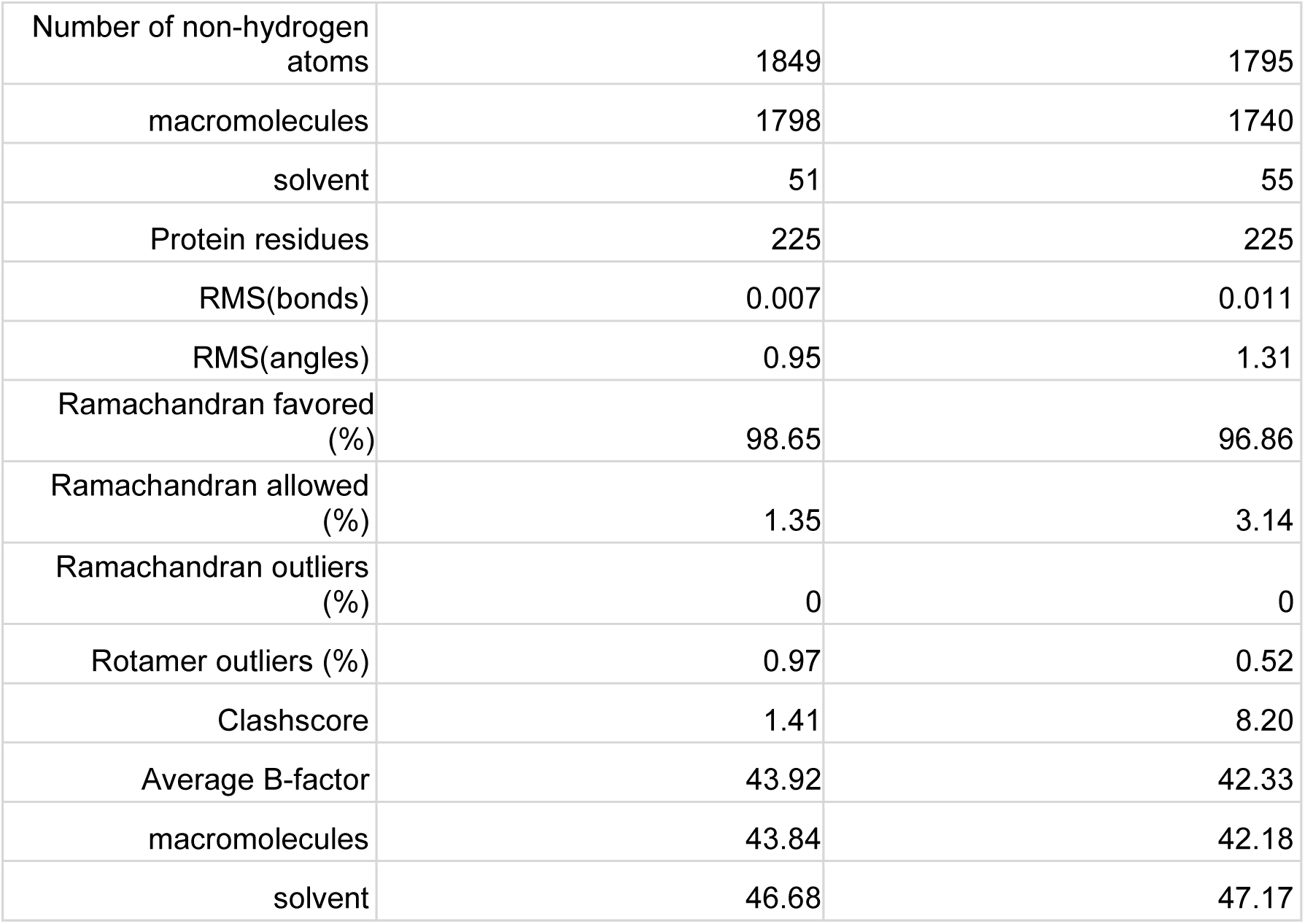
Crystallographic data and structural refinement statistics.

|  | **REP24** | **REP24** |
| --- | --- | --- |
| Data collection source | SSRL, 12-1 | ESRF, ID29 |
| Data collection method | Merged rotation series from multiple crystals | Serial crystallography |
| Data collection temperature (°C) | 21 | 21 |
| PaN Fixed target format | Open chip loading | Film-on-film |
| Wavelength (keV) | 12.4 | 11.6 |
| Resolution range (Å) | 39.15 - 1.6 (1.66 - 1.6) | 39.14 - 1.8 (1.9 - 1.8) |
| Space group | P 32 2 1 | P 32 2 1 |
| Unit cell (a, b, c, $\alpha$, $\beta$, $\gamma$) | 45.209 Å, 45.209 Å, 183.917 Å, 90°, 90°, 120° | 45.2 Å, 45.2 Å, 183.2 Å, 90°, 90°, 120° |
| Total reflections | 656553 (62888) | 9622135 (400928) |
| Unique reflections | 40080 (3899) | 21055 (2948) |
| Multiplicity | 16.4 (16.1) | 457 (136) |
| Completeness (%) | 95.66 (81.74) | 99.75 (99.73) |
| Mean I/sigma(I) | 5.26 (0.14) | 5.1 (0.2) |
| Wilson B-factor | 32.73 | 34.07 |
| R-merge | 0.2345 | -- |
| R-split | -- | 10.9 |
| R-work | 0.2084 | 0.2506 |
| R-free | 0.2264 | 0.3169 |
| Number of non-hydrogen atoms | 1849 | 1795 |
| macromolecules | 1798 | 1740 |
| solvent | 51 | 55 |
| Protein residues | 225 | 225 |
| RMS(bonds) | 0.007 | 0.011 |
| RMS(angles) | 0.95 | 1.31 |
| Ramachandran favored (%) | 98.65 | 96.86 |
| Ramachandran allowed (%) | 1.35 | 3.14 |
| Ramachandran outliers (%) | 0 | 0 |
| Rotamer outliers (%) | 0.97 | 0.52 |
| Clashscore | 1.41 | 8.20 |
| Average B-factor | 43.92 | 42.33 |
| macromolecules | 43.84 | 42.18 |
| solvent | 46.68 | 47.17 |

## Supporting information

Supplementary File: Fabrication details

Supplementary Movie: PaN fixed-target data collection at ESRF

## Acknowledgement

This work was performed, in part, under the auspices of the U.S. Department of Energy by Lawrence Livermore National Laboratory under Contract No. DE-AC52-07NA27344. This work was supported by the National Institute of Health (NIH) Grant No. R01GM117342 (NIGMS). The contents of this publication are solely the responsibility of the authors and do not necessarily represent the official views of NIGMS or the NIH. We acknowledge beamtime provision at the European Synchrotron Radiation Facility (ESRF) granted from ID29 BAG proposals MX2578 and would like to thank the ESRF - EMBL Joint Structural Biology Group for continuous support. We acknowledge the use of the Stanford Synchrotron Radiation Lightsource (SSRL) and Linac Coherent Light Source at the SLAC National Accelerator Laboratory supported by the US Department of Energy, Office of Science, Office of Basic Energy Sciences under contract No. DE-AC02-76SF00515. The SSRL Structural Molecular Biology Program is supported by the DOE Office of Biological and Environmental Research and by the National Institutes of Health, National Institute of General Medical Sciences (P30GM133894). Sabine Botha would like to acknowledge support by the National Science Foundation Bio Directorate under midscale research infrastructure Grants 2153503 and 1935994. This research was supported in part by an appointment to the NNSA Minority Serving Institutions Internship Program (Chandraki Chatterjee), sponsored by the U.S. Department of Energy and administered by the Oak Ridge Institute for Science and Education. Crystallization assistance at the National Crystallization Center at UB HWI was supported through NIH grant R24GM141256.

## Supplementary Material

### Parylene coater handling procedure

The following procedure is followed to obtain nearly uniform Parylene-N deposition on the 6” wafer. SCS parylene coater is utilized to fabricate the free-standing PaN films.

1. Inspect and clean the Cold Trap

- Check for parylene build-up.
- If build-up is present, clean the cold trap thoroughly.
- After cleaning, coat all surfaces with diluted Micro-90 to prevent adhesion of deposited material.
2. Prepare the Chamber

- Remove the chamber lid and place it upside down on a clean surface. Ensure the O-ring does not touch any surface.
- Using Scotch tape, remove debris from the chamber lid’s O-ring. Confirm the O-ring is clean and intact.
- Inspect the port holes inside the chamber and ensure they are not obstructed by parylene.
3. Load silicon wafers

- Place the silicon wafers in the center of the chamber.
4. Reassemble the Chamber

- Confirm that the O-ring seating area is free of debris.
- Carefully reposition the chamber lid, ensuring a flush seal with no visible gaps.
5. Load Parylene Dimer

- Prepare a boat-shaped aluminum foil container.
- Weigh out the desired amount of dimer and place it into the boat.
- Open both the dimer chamber and the secondary chamber.

◦ Inspect and clean O-rings as necessary.
- Load the dimer-containing boat into the chamber.
- Close all chamber doors securely.
6. Install the Cold Trap

- Ensure the O-ring on the cold trap is clean before placing it into position.
7. Power On System

- If previously pressed, depress the EMO button.
- Switch on main power and allow the system to reset.
8. Confirm Deposition Parameters

- Input the appropriate deposition parameters for the experiment.
9. Initiate Vacuum

- Turn on the vacuum system and allow to run for 15 minutes.
10. Cool the Cold Trap

- Switch on the chiller and wait 30–60 minutes until the cold trap is sufficiently cooled or chamber pressure drops below 30 Torr.
- Ensure vacuum remains ON throughout.
11. Begin Heating Sequence

- Turn on the furnace and chamber gauge heater.
- Turn on the vaporizer heater.
12. Begin Deposition

- Press the green start button.

◦ Confirm sample rotation visually through the chamber window.
- The system will:

◦ Heat the furnace and gauge heaters to operating temperature.
◦ Maintain vacuum until base pressure (∼17 Torr) is reached.
◦ Ramp up the vaporizer heater, causing dimer vaporization.
◦ Pyrolyze the dimer into monomer gas, which deposits within the room temperature chamber.
- Monitor deposition:

◦ Chamber pressure increases during deposition.
◦ Once complete, pressure declines.
◦ Upon reaching base pressure again, heaters will shut off automatically.
13. End Deposition

- Press the green stop button.
- Turn OFF the furnace and chamber gauge heaters.
- Turn OFF the vaporizer heater.
- Keep the vacuum and chiller ON.
14. Cool Down System

- Allow the system to cool until the furnace temperature is <1000 °C (acceptable up to 1500 °C).
- Cooling may take up to 3 hours.
- If water condensation appears on the cold trap lid, wipe dry with a lint-free cloth.
15. Final Shutdown

- Turn OFF the chiller.
- Vent the chamber—this should take only a few seconds.
16. Disassemble System

- Carefully remove the cold trap:

◦ Avoid bending the tube, as cold glass is prone to cracking.
◦ If water vapor condenses into ice on a room-temperature trap, allow it to melt before reuse.
- Remove the chamber lid. This may require gentle pressure.
- Retrieve the silicon wafer (if used).
17. Re-seal Chamber

- Clean the chamber lid O-ring again with Scotch tape.
- Reinstall the chamber lid.
- Set the vacuum to HOLD mode or return the chamber to atmospheric conditions as standard practice.
18. Final Steps

- Remove or save the aluminum dimer boat for reuse or disposal.
- Press the EMO button to shut down the system.
- After wafer removal, retrieve ultra-thin free-standing PaN films using tweezers.

◦ Use two tweezers to prevent folding or damage due to film fragility.

**Fig. S1:**
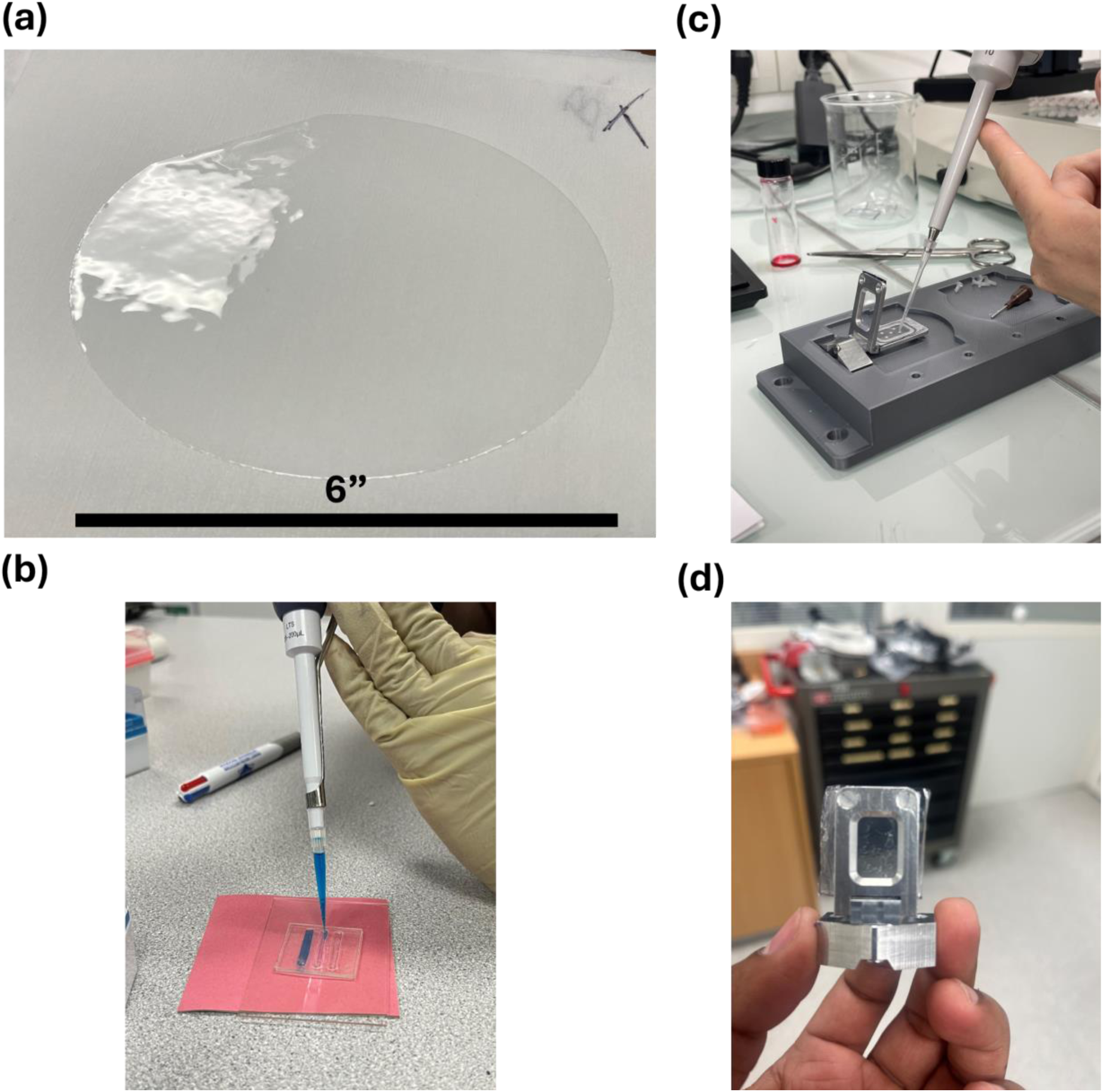
a) Large area free-standing PaN film. These films are cut to size to make different types of PaN fixed-target sample platforms; b) Closed-chip PaN FT with independent windows; c) REP-24 microcrystals sample loading on film-on-film PaN FT and held in place to the actuators using SOS metal holder^49^; d) Film-on-film PaN FT loaded with REP-24 wild type protein microcrystals.

#### Parameters utilized for screening printing to fabricate closed chip PaN fixed-target platforms

**Fig. S2.**
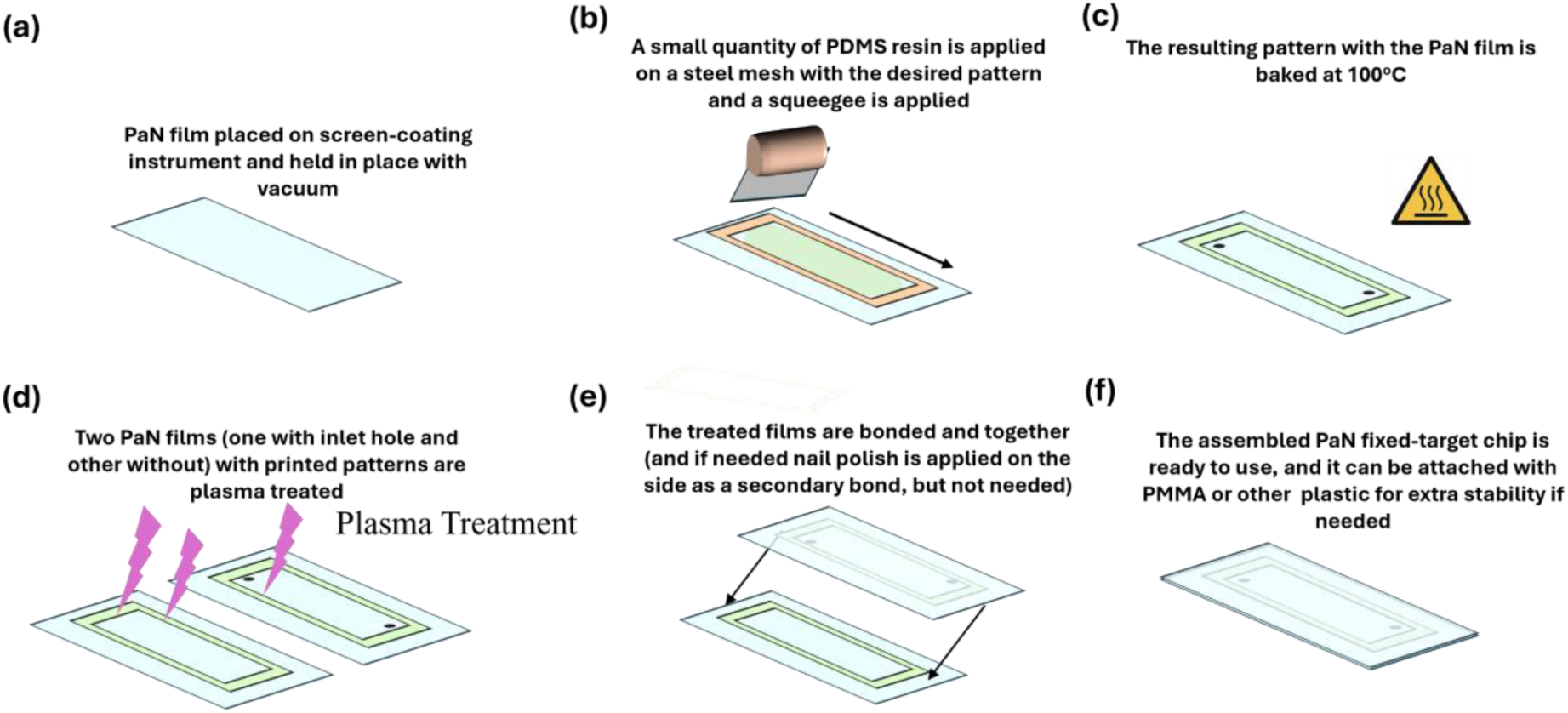
Systematic Screen printing methodology of making closed-chip PaN FT. a) The PaN film is cut to size, and placed on the screen-coating instrument, and held in place with vacuum; b) A stell-mesh with the desired pattern is placed on the film, and a small quantity of PDMS resin is introduced and the squeegee is applied; c) The resulting pattern with the PaN film is baked at 100°C; d) Two patterned films are plasma treated; e) The plasma treated films are placed one-upon-another and bonded; f) The ready to use assembled devices have holes at the desired locations for crystal slurry introduction.

For screen printing, the relationship between screen parameters and ink rheological characteristics is of outmost importance. If an ink is too viscous, no matter how large the mesh opening it will not effectively pass through the mesh to achieve a successful print. However, if it is too thin, the ink will likely bleed out from the desired pattern when deposited, which is an undesirable outcome. The ideal ink is one with reasonably high viscosity and with the ability to shear-thin. This ensures they can keep their shape while in rest but also flow when exposed to large pressures like that of going through a screen. Here the ink is a PDMS mixture (10:1 ratio of base to curing agent). Mesh count (MC), which is the measure of how many threads cross each other in a square inch of screen has a large contribution to feature printability. Also depicted with its metric units, threads/cm, high mesh counts usually lead to fine feature printing when paired with the correct wire thickness and emulsion thickness. The emulsion is the coating over the mesh that masks off any areas you do not want the ink to transfer through. It also greatly influences the quality of the print as it helps with detail resolution and ink transfer. In this work, we can see how the pairing of different MC screen openings and emulsion thicknesses led to different print thicknesses when paired with either a thick or thin ink of PDMS. Wire diameter is also an important factor in final print thickness, however, it was not varied within MCs in this study and therefore not a significant factor in this work.

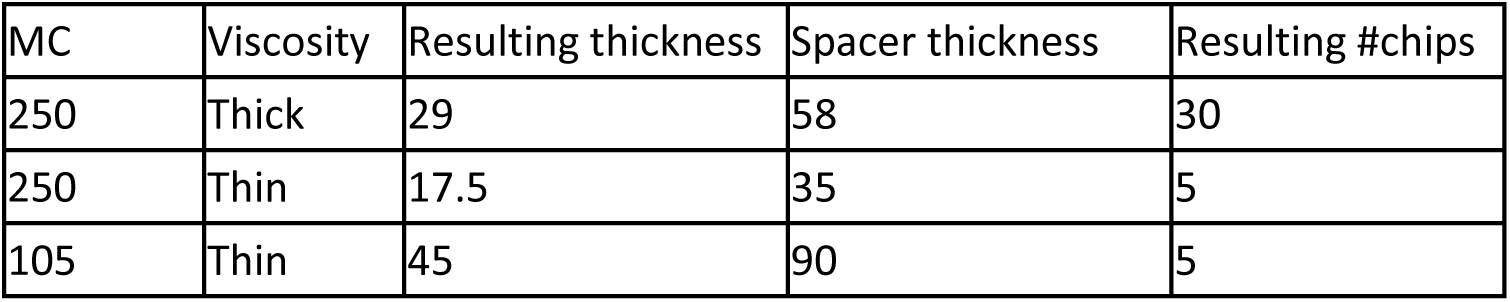

Screen Parameters

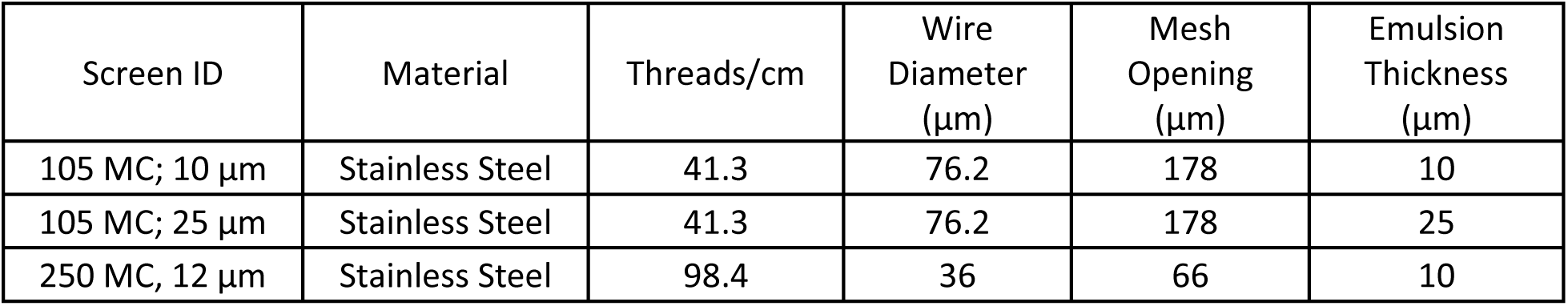

Combination Mesh + Ink Viscosity on resulting thickness

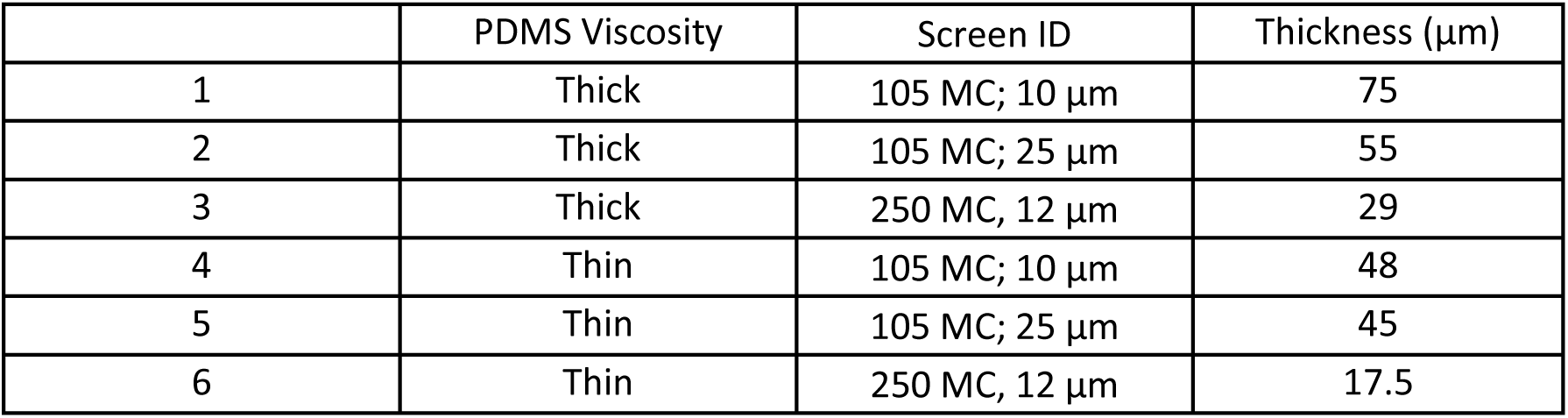

### X-ray Fluence calculation

Based on the x-ray beam spot size of ∼3 µm, the spot area is calculated to be 7.07 × 10^-12^ m^2^.. Converting 9.6 keV photon energy in Joules, we obtain as 1.54 × 10^-15^ J. Number of photons per pulse is pulse energy/photon energy, i.e., 1.04 × 10^12^. Photon fluence (i.e., photons per unit area) for 3 µm spot size, is 1.47 × 10^19^ photons/cm^2^. Energy fluence is determined as E/A (where E is the pulse energy, and A is the spot area), which is 2.26 × 10^4^ J/cm^2^.

**Fig S3.**
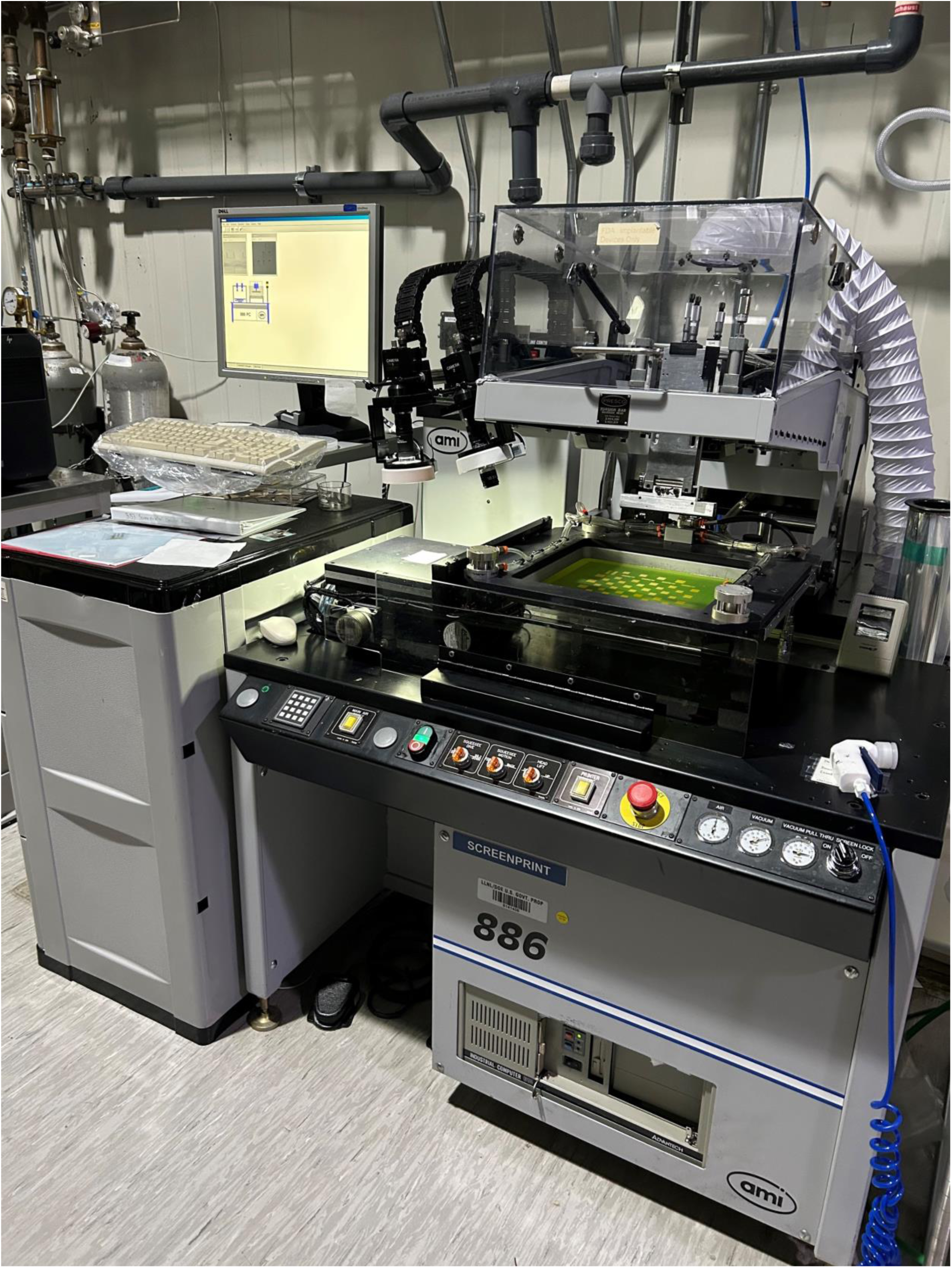
Automated vacuum-holder based screen printer utilized for fabricating closed chip PaN FT.

