## Supplementary File: Fabrication details for "Novel polymer fixed-target microfluidic platforms with an ultra-thin moisture barrier for serial macromolecular crystallography"

LLNL-JRNL-2007768

1. Biosciences and Biotechnology Division, Physical and Life Sciences Directorate, Lawrence Livermore National Laboratory, Livermore, USA; 2. Materials Engineering Division, Engineering Directorate, Lawrence Livermore National Laboratory, Livermore, USA; 3. ESRF- The European Synchrotron, Grenoble, France; 4. Department of Applied Physics, Graduate School of Engineering, Nagoya University; Nagoya, Japan; 5. Department of Physics, Arizona State University, Arizona, USA; 6. Department of Chemical Engineering, University of California, Davis, USA; 7. Department of Chemistry, The State University of New York at Buffalo, Amherst, USA; 8. Linac Coherent Light Source, SLAC National Accelerator Laboratory, Menlo Park, USA; 9. Materials Science Division, Physical and Life Sciences Directorate, Lawrence Livermore National Laboratory, Livermore, USA; 10. Stanford Synchrotron Radiation Light Source, SLAC National Accelerator Laboratory, Menlo Park, USA; 11. Hauptman-Woodward Medical Research Institute, Buffalo, USA; 12. European Molecular Biology Laboratory, Grenoble, France; 13. Department of Biochemistry and Molecular Medicine, School of Medicine, University of California, Davis, USA.

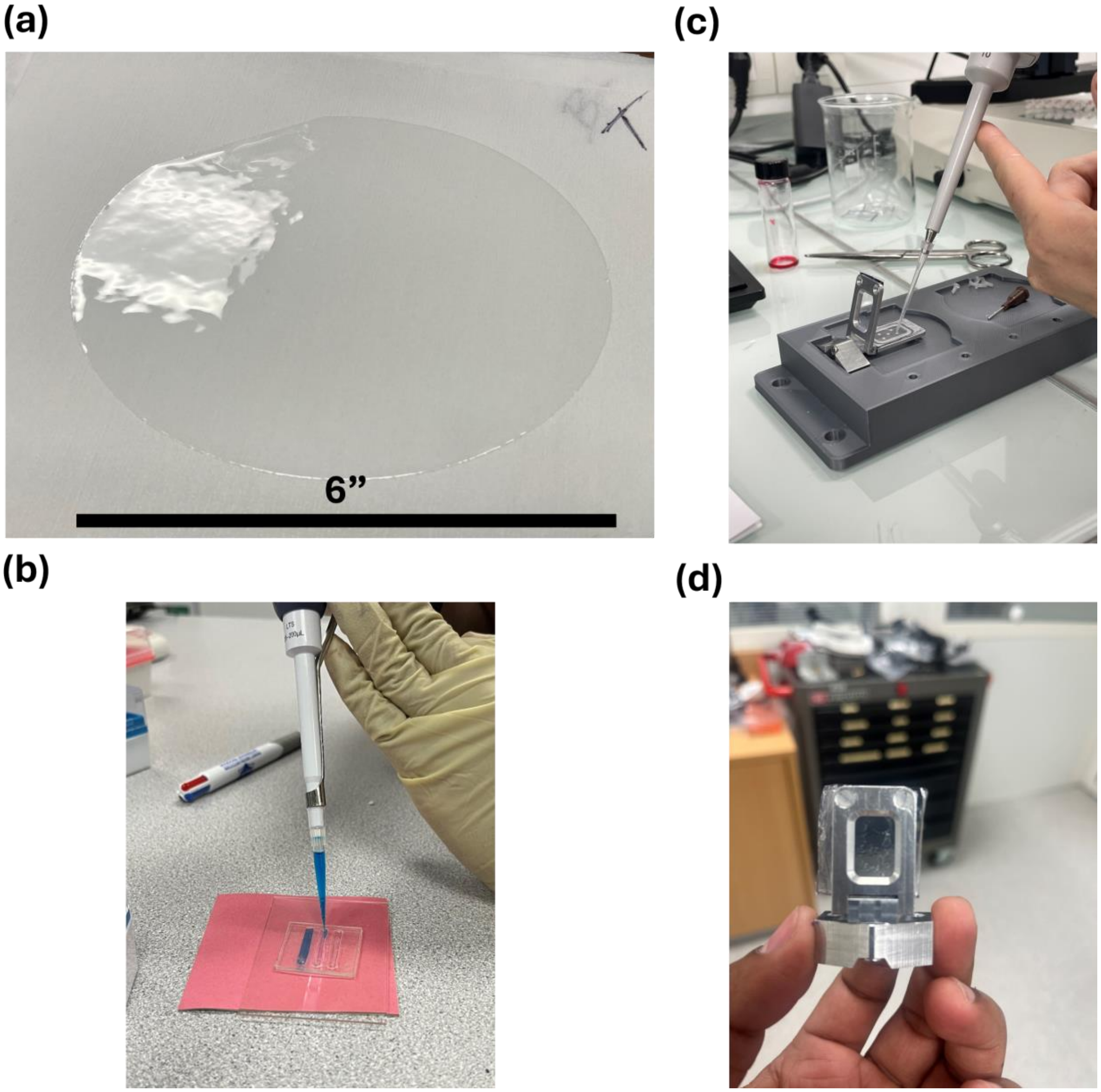


Fig. S1: a) Large area free-standing PaN film. These films are cut to size to make different types of PaN fixed-target sample platforms; b) Closed-chip PaN FT with independent windows; c) REP-24 microcrystals sample loading on film-on-film PaN FT and held in place to the actuators using SOS metal holder[49]; d) Film-on-film PaN FT loaded with REP-24 wild type protein microcrystals.

**Parameters utilized for screening printing to fabricate closed chip PaN fixed-target platforms:**

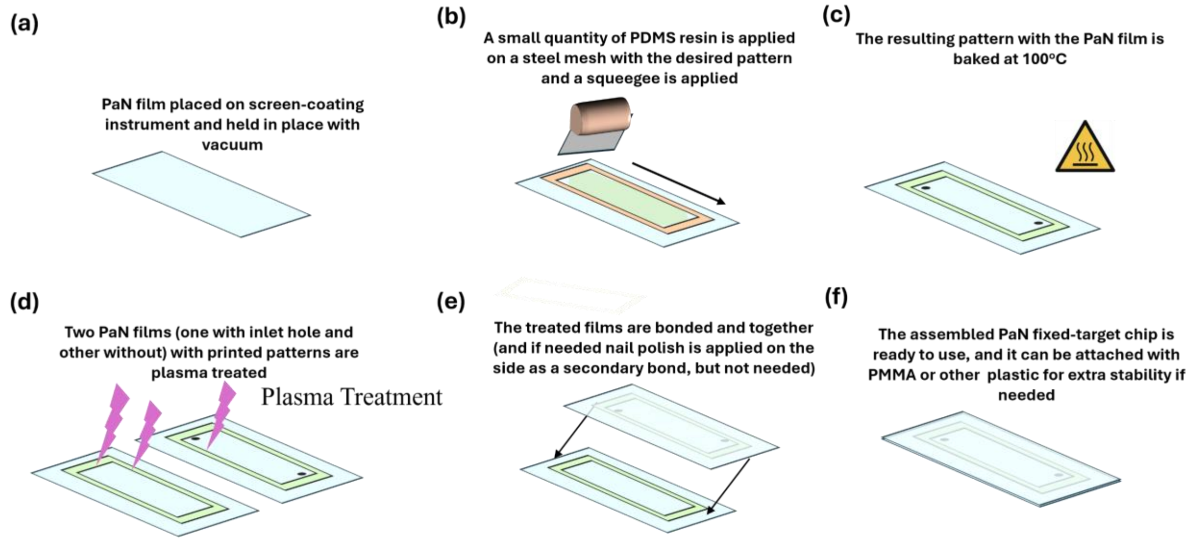


Fig. S2. Systematic Screen printing methodology of making closed-chip PaN FT. a) The PaN film is cut to size, and placed on the screen-coating instrument, and held in place with vacuum; b) A stell-mesh with the desired pattern is placed on the film, and a small quantity of PDMS resin is introduced and the squeegee is applied; c) The resulting pattern with the PaN film is baked at 100°C; d) Two patterned films are plasma treated; e) The plasma treated films are placed one-upon-another and bonded; f) The ready to use assembled devices have holes at the desired locations for crystal slurry introduction.

For screen printing, the relationship between screen parameters and ink rheological characteristics is of outmost importance. If an ink is too viscous, no matter how large the mesh opening it will not effectively pass through the mesh to achieve a successful print. However, if it is too thin, the ink will likely bleed out from the desired pattern when deposited, which is an undesirable outcome. The ideal ink is one with reasonably high viscosity and with the ability to shear-thin. This ensures they can keep their shape while in rest but also flow when exposed to large pressures like that of going through a screen. Here the ink is a PDMS mixture (10:1 ratio of base to curing agent). Mesh count (MC), which is the measure of how many threads cross each other in a square inch of screen has a large contribution to feature printability. Also depicted with its metric units, threads/cm, high mesh counts usually lead to fine feature printing when paired with the correct wire thickness and emulsion thickness. The emulsion is the coating over the mesh that masks off any areas you do not want the ink to transfer through. It also greatly influences the quality of the print as it helps with detail resolution and ink transfer. In this work, we can see how the pairing of different MC screen openings and emulsion thicknesses led to different print thicknesses when paired with either a thick or thin ink of PDMS. Wire diameter is also an important factor in final print thickness, however, it was not varied within MCs in this study and therefore not a significant factor in this work.

| MC | Viscosity | Resulting thickness | Spacer thickness | Resulting #chips |
|---|---|---|---|---|
| 250 | Thick | 29 | 58 | 30 |
| 250 | Thin | 17.5 | 35 | 5 |
| 105 | Thin | 45 | 90 | 5 |

Screen Parameters

| Screen ID | Material | Threads/cm | Wire Diameter (µm) | Mesh Opening (µm) | Emulsion Thickness (µm) |
|---|---|---|---|---|---|
| 105 MC; 10 µm | Stainless Steel | 41.3 | 76.2 | 178 | 10 |
| 105 MC; 25 µm | Stainless Steel | 41.3 | 76.2 | 178 | 25 |
| 250 MC, 12 µm | Stainless Steel | 98.4 | 36 | 66 | 10 |

Combination Mesh + Ink Viscosity on resulting thickness

| | PDMS Viscosity | Screen ID | Thickness (µm) |
|---|---|---|---|
| 1 | Thick | 105 MC; 10 µm | 75 |
| 2 | Thick | 105 MC; 25 µm | 55 |
| 3 | Thick | 250 MC, 12 µm | 29 |
| 4 | Thin | 105 MC; 10 µm | 48 |
| 5 | Thin | 105 MC; 25 µm | 45 |
| 6 | Thin | 250 MC, 12 µm | 17.5 |

<u>X-ray Fluence calculation:</u>

Based on the x-ray beam spot size of ~3 µm, the spot area is calculated to be 7.07 x $10^{-12}$ $m^2$.. Converting 9.6 keV photon energy in Joules, we obtain as 1.54 x $10^{-15}$ J. Number of photons per pulse is pulse energy/photon energy, i.e., 1.04 x $10^{12}$. Photon fluence (i.e., photons per unit area) for 3 µm spot size, is 1.47 x $10^{19}$ photons/$cm^2$. Energy fluence is determined as E/A (where E is the pulse energy, and A is the spot area), which is 2.26 x $10^4$ J/$cm^2$.

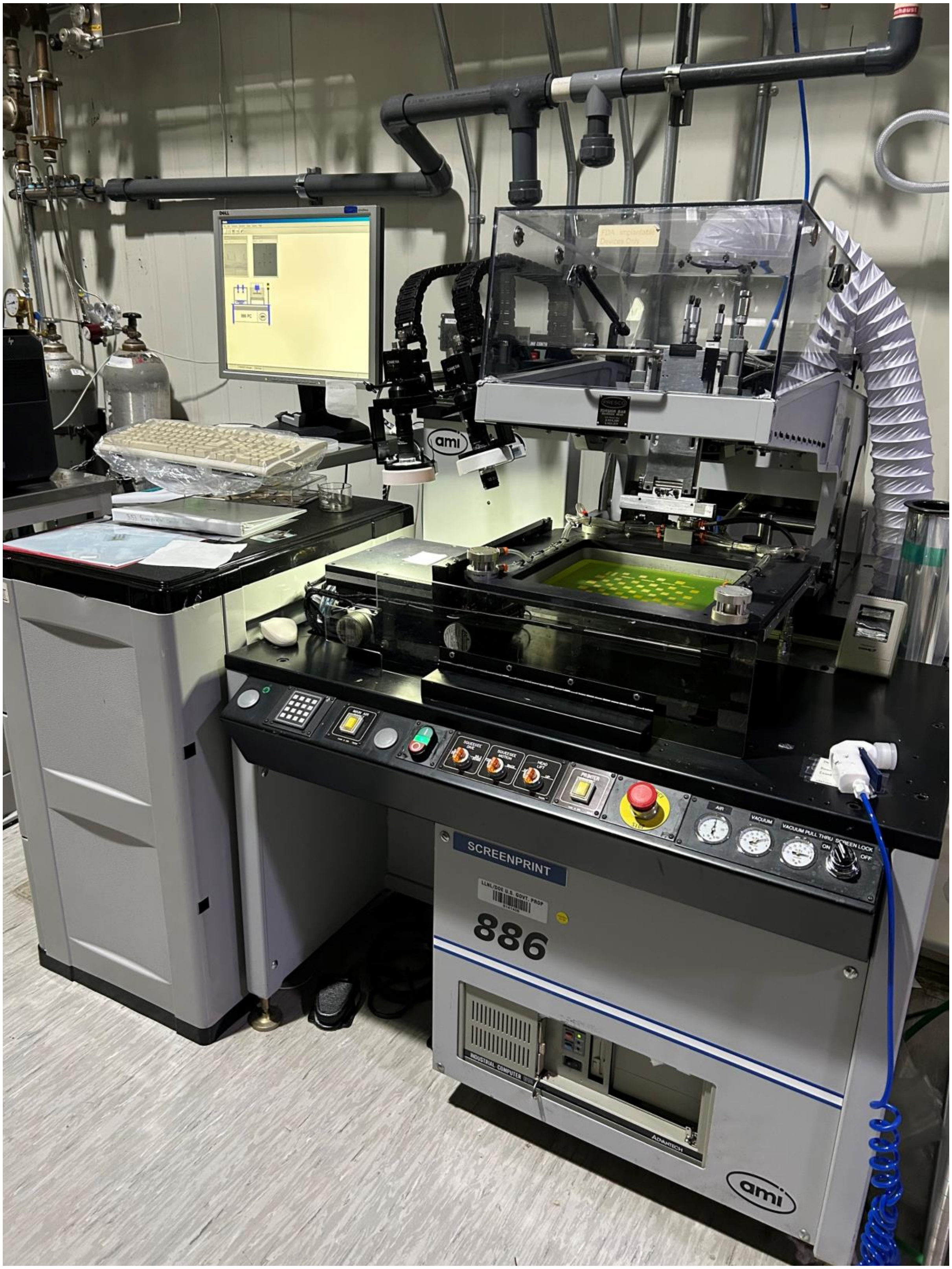


Fig S3. Automated vacuum-holder based screen printer utilized for fabricating closed chip PaN FT.
